# Ingestion of the potent neurotoxin epibatidine does not compromise locomotion or behavior in the poison frog *Epipedobates tricolor*

**DOI:** 10.64898/2026.06.10.731460

**Authors:** Adriana M. Jeckel, Sophie J. Draper, Benjamin P. Deters, Rachel B. Weinberg, Neil D. Tsutsui, Rebecca D. Tarvin

## Abstract

Most organisms have evolved mechanisms to reduce the negative effects of toxins in their diet. Some animals that are toxin specialists, such as dendrobatid poison frogs, have amino acid substitutions in proteins targeted by the toxins that prevent or limit the toxin’s ability to bind and exert its bioactive effects. The Phantasmal poison frog, *Epipedobates tricolor*, has amino acid substitutions in its neuronal nicotinic acetylcholine receptors that were previously shown to provide resistance to the highly potent neurotoxin epibatidine *in vitro.* However, it is unclear whether *E. tricolor* resists the physiological effects of epibatidine *in vivo*. To investigate this, we examined the effects of epibatidine exposure on the locomotion and behavior of *E. tricolor*. We performed whole-animal performance assays and behavioral evaluations at multiple time points following the administration of high yet biologically relevant levels of epibatidine. These assays were followed by alkaloid quantification to track chemical concentrations in the skin. *Epipedobates tricolor* exhibited similar locomotor performance and behavior at both epibatidine doses compared to controls and regardless of the quantity of alkaloid accumulated into the skin. However, we observed an impact of low-percentage ethanol solutions on behavior when compared to water controls, as well as general impacts of handling stress (regardless of the administered solution type), which should be considered in future experimental designs. Overall, we demonstrate that *E. tricolor* likely avoids a physiological fitness trade-off between toxin ingestion and the defensive benefits of epibatidine sequestration. Our study suggests that the ability to ingest toxins involves multi-faceted resistance mechanisms.

**Highlights:**

– Whole-animal performance assay shows no impact of epibatidine on frog behavior
– *Epipedobates tricolor* is 20–5000 times more resistant to epibatidine than mice
– Assays demonstrate that frogs are sensitive to 6% ethanol and handling stress

## 1. INTRODUCTION

Across diverse taxa, consumers possess strategies to mitigate the effects of dietary toxins. These mechanisms range from behavioral avoidance to metabolic detoxification or intricate molecular modifications of biological targets (Tarvin et al., 2023). Even with some resistance, ingesting these toxins likely still carries some cost, which may come in the form of reduced energy intake due to behavioral choices (*e.g.*, choosing not to consume available but toxic items; Köhler et al., 2012; Lerch-Henning and Nicolson, 2013; Ramírez-Castañeda et al., 2025) or physiological constraints (*e.g.*, reduced gut transit and sugar assimilation due to the presence of toxins; Tadmor-Melamed et al., 2004), allocation costs (energy expenditure to detoxify toxins at the cost of growth, fecundity, or immune defense; Agrawal et al., 2021; Camara, 1997; Goetz et al., 2018; Manson and Thomson, 2009; Speed and Ruxton, 2014), or functional constraints (*e.g.*, trade-off between protein function and toxin insensitivity; Feldman et al., 2012; Geffeney et al., 2002, 2005).

Dietary toxins often target membrane receptors and ion channels, directly interfering with neuronal and muscular functions. Some animals that specialize on toxic food sources have target-site resistance (TSR), in which amino acid substitutions in the toxin binding site prevent or limit the ability of the toxin to bind its molecular target and help subvert its primary adverse effects. For example, several *Thamnophis* snake species prey upon *Taricha* newts, which possess the potent neurotoxin tetrodotoxin (TTX).

Although TTX normally blocks voltage-gated sodium channels, amino acid substitutions in the snakes’ skeletal muscle sodium channel reduce the binding site’s affinity for TTX, conferring TSR (Geffeney et al., 2005). Despite this resistance, *Thamnophis sirtalis* snakes remain incapacitated for thirty minutes to several hours after ingesting the toxin, suggesting that TSR may be insufficient to counteract the full impact of TTX exposure (Williams et al., 2002). Further, some populations of *T. sirtalis* possess specific amino acid substitutions in the sodium channel outer pore that confer increased whole-animal resistance but reduce channel excitability, which lowers muscle performance more generally (Brodie III and Brodie Jr., 1999; Feldman et al., 2012; Geffeney et al., 2002, 2005; Hague et al., 2018). Despite the apparent drawbacks of toxin consumption, many animals choose to consume toxins, likely because of related fitness benefits, such as reduced competition for prey or accumulation of toxins for chemical defense.

While TSR is one mechanism allowing for toxin consumption, organisms that have evolved the ability to ingest and accumulate dietary toxins (*i.e.*, sequestration) may require multiple forms of resistance to offset the physiological costs imposed by constant and high levels of toxin exposure. Monarch butterflies (*Danaus plexippus*), for instance, have a TSR mechanism that enables them to consume milkweed plants containing cardiac glycosides, which are Na^+^/K^+^-ATPase inhibitors, and store them for protection against predators (Brower et al., 1968; Yatime et al., 2011). Despite having TSR, monarch larvae are still negatively affected by very high concentrations of these compounds in the leaf latex, resulting in symptoms such as temporary muscular rigidity, fixed posture, and reduced responsiveness (Agrawal et al., 2021; Zalucki et al., 2001). They avoid feeding on the latex by cutting the leaf veins and draining the latex from the leaf prior to feeding (Dussourd, 1999; Dussourd and Eisner, 1987). By employing an additional resistance mechanism, they are able to still feed on the compounds and sequester them, while avoiding the costs of the high concentrations.

Poison frogs (family Dendrobatidae) of Central and South American rainforests are famous for their ability to ingest and sequester dietary alkaloids for defense against predators and pathogens (Daly et al., 1994b, 1994a; Saporito et al., 2012). These frogs can sequester a wide range of alkaloids: at least 1,200 alkaloids across 26 classes have been described in the skin of wild-caught poison frogs (Daly et al., 2005; Saporito et al., 2012). This high diversity alone suggests that resistance involves multiple mechanisms, yet it is not comprehensively characterized in this group. The best known example is the TSR in the β subunit of neuronal nicotinic acetylcholine receptors (nAChRs), which provides resistance to the highly toxic alkaloid epibatidine (Tarvin et al., 2017).

Epibatidine, a potent nAChR agonist with 20 times the affinity of nicotine and 200 times the analgesic potency of morphine (Fig. 1A, Badio and Daly, 1994), was first detected in the dendrobatid poison frog *Epipedobates anthonyi*, from southwestern Ecuador (López-Hervas et al., 2024; Spande et al., 1992). *Epipedobates* is a small genus in the family Dendrobatidae containing seven recognized species found on the western side of the Andes from northern Peru to central Colombia, with extensive variation in color pattern and in skin alkaloid profiles (Daly et al., 2000; López-Hervas et al., 2024; Tarvin et al., 2017b). An amino acid substitution at position S108C in the nAChR β2 subunit present in *Epipedobates* and other poison frog species confers resistance to epibatidine but may impose costs on the organism because the toxin shares a binding site with the endogenous ligand, acetylcholine. While additional substitutions in the β2 subunit appear to compensate for reduced acetylcholine sensitivity in *Epipedobates* and *Ameerega* poison frogs, *Oophaga* species lack compensatory substitutions (Tarvin et al., 2017a). Tarvin et al. (2016) identified additional putative TSR-conferring substitutions in the voltage-gated sodium channel NaV1.4 using computational modeling; although one of these mutations may provide resistance to the neurotoxin batrachotoxin (Abderemane-Ali et al., 2021; Wang and Wang, 2017), the contribution of most of these substitutions to alkaloid resistance has yet to be tested. Other mechanisms may contribute to epibatidine resistance in *Epipedobates* poison frogs, including avoidance behavior, metabolism, transport proteins, and other alternative targets (Alvarez-Buylla et al., 2023; Caty et al., 2019; Douglas et al., 2022; Jeckel et al., 2026; Tarvin et al., 2024). A plasma globulin-like protein in a few species of poison frogs has been characterized as binding to several alkaloids; however, in *Epipedobates tricolor*, the tested protein does not react with epibatidine (Alvarez-Buylla et al., 2023). In addition, in controlled laboratory feeding experiments, *Epipedobates anthonyi* sequestered 6% of the total ingested alkaloids (Waters et al., 2023). This suggests that most of the ingested epibatidine does not reach the skin glands and indicates the presence of alternative pathways for ingested alkaloids. While these data suggest that some poison frogs have resistance to alkaloids, the *in vivo* costs of toxin exposure remain unknown.

**Figure 1.**
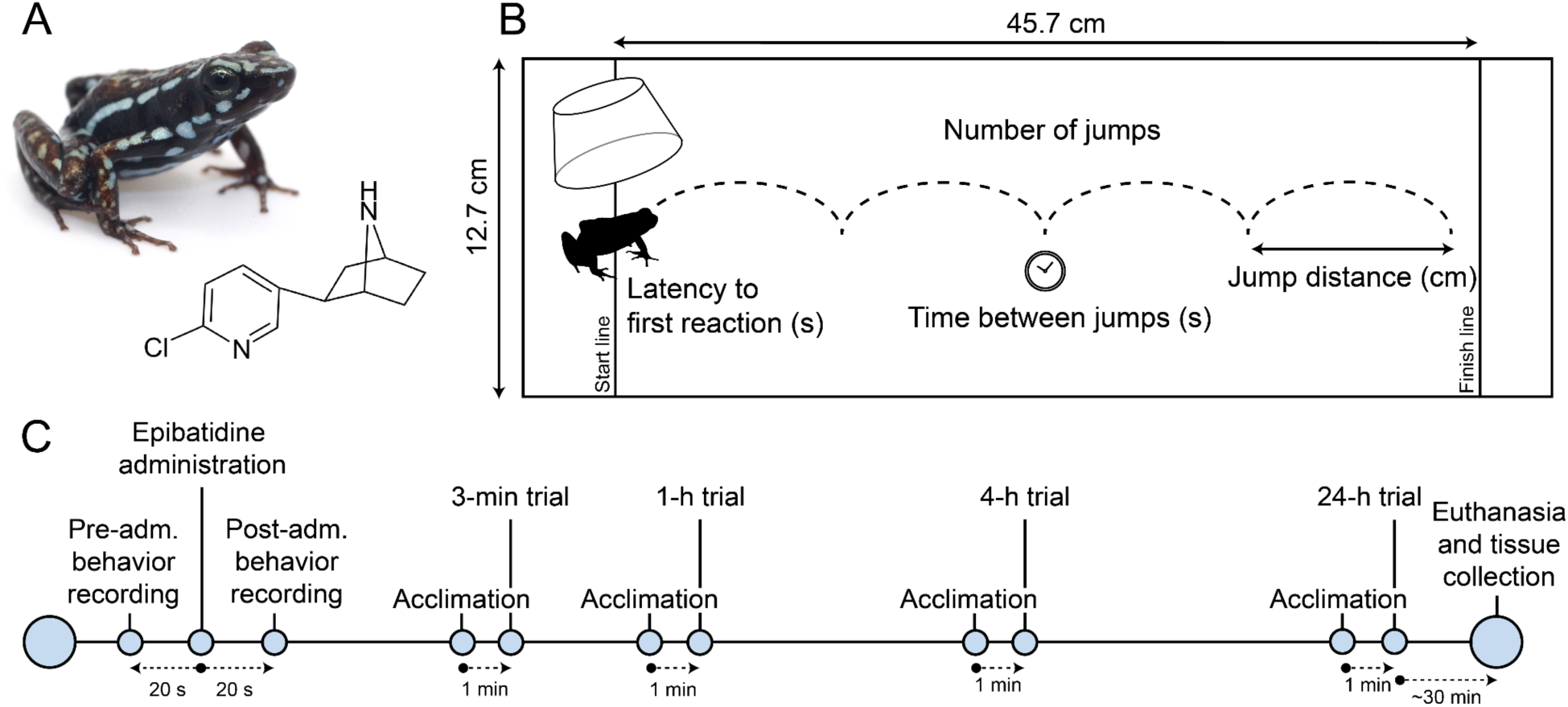
Whole-animal performance assays. A) Captive bred specimen of *Epipedobates tricolor* “Cielito” and chemical structure of epibatidine. B) Schematic image of the arena and the parameters analyzed. C) Timeline of the whole-animal performance assays per frog.

The main goal of this study was to test whether epibatidine exposure affects locomotion or behavior of the poison frog *Epipedobates tricolor in vivo*. We performed whole-animal performance assays to evaluate the effect of different doses of epibatidine at different time points following alkaloid administration. By pairing performance assays with alkaloid quantification, we aimed to evaluate whether other forms of resistance may also play a role in offsetting the physiological cost of exposure to epibatidine in *E. tricolor*.

## 2. MATERIAL AND METHODS

### 2.1 Animals

We selected 33 individuals (N = 10 per epibatidine treatment, N = 8 for ethanol control group, and N = 5 for water control group) of *Epipedobates tricolor* “Cielito” (Fig. 1A) ranging in mass from 0.5–0.8 g from a breeding colony housed at University of California, Berkeley under IACUC protocol AUP-2023-03-16154-1, originally sourced from Indoor Ecosystems, LLC (Whitehouse, OH, USA). On the day of the experiment, we weighed each frog, photographed their dorsal and ventral pattern for identification, and kept them in 30 x 17 x 10 cm terraria in groups of three with soil and *Sphagnum* sp. moss as substrate. Throughout the experiment, animals were kept in a 12 h:12 h light:dark cycle, with humidity >80% and temperature 23–25°C. We fed the frogs daily with *Drosophila melanogaster* fruit flies dusted with vitamin powder and carotenoids (Calcium Plus, Vitamin A Plus, and SuperPig, Repashy Ventures Inc, Oceanside, CA, USA).

### 2.2 Toxin feeding

We used four experimental groups: water control, ethanol control, 2-µg epibatidine, and 6-µg epibatidine. The 2-µg treatment was based on the estimated average daily consumption of alkaloids in nature. On the basis of stomach content of dietary ants in wild caught adult *E. tricolor* (data not published), we estimated a maximum consumption of 11 ants per day, which would total 3.3 µg of alkaloids per day, based on the average amount of alkaloids in ants (Jones et al., 1996; Jones and Blum, 1982, also see Jeckel et al., 2022). We corrected for the difference in average body mass between adult *E. tricolor* (∼0.9 g) and the individuals used in our study (0.6 ± 0.1 g), resulting in a full daily dosage of 2.5 μg of total alkaloid, which we rounded down to 2.0 μg to have a more conservative quantity of alkaloids for the lower dose treatment.

The 6-µg treatment was selected as a relatively high dose that was below the limit observed anecdotally to cause lethargy and paralysis (10 µg/day *sensu* Waters et al., 2023). Epibatidine is one of the most potent alkaloids found in poison frogs (Santos et al., 2016), yet it is found in very low concentrations in nature (Daly et al., 2000; Spande et al., 1992; Tarvin et al., 2024), and thus 6.0 µg epibatidine is expected to represent a physiologically relevant challenge. (±)-Epibatidine was purchased commercially (CAS: 162885-01-0, Sigma-Aldrich, Darmstadt, Germany) and dissolved in a 2 or 6% ethanol solution (2-µg treatment: 4.0 µL of 0.5 µg/µL epibatidine in 2% ethanol; 6-µg treatment: 4.0 µL of 1.5 µg/µL epibatidine in 6% ethanol). We decided to feed the ethanol control group frogs a 6% ethanol solution with 0 µg epibatidine (rather than 2% ethanol), comparable to the 6-µg treatment, to best capture the effect of ethanol on the frogs. We also included a water control group, where we administered deionized water remineralized with R/O Rx according to manufacturer’s instructions (Josh’s Frogs, Owosso, MI, USA). We administered 4.0 µL of solution to each individual by opening their mouth with a disposable plastic pipette and inserting the tip of the micropipette into the oral cavity to deliver the alkaloid solution (following Jeckel et al., 2022).

### 2.3 Mouth-gaping behavior

Mouth-gaping behavior is a common sign of distress in amphibians (Hurlbert, 1970; Taylor et al., 2021), and it is unusual for *E. tricolor* to open its mouth, except for eating (personal observation). Approximately 15 minutes before the oral administration and the performance assays, we collected all the frogs from their terraria and kept them in individual transparent cups (7.5 cm diameter) in a dark environment to limit additional stress. We recorded a 20-second video of each frog immediately before and immediately after solution administration using an iPhone 13 [iOS 26.3]. We considered a mouth-gaping event every time the frog opened and closed its mouth, independently of how long the mouth remained open. To avoid external factors of stress that could bias the mouth-gape count (*e.g.*, escaping from the arena), we excluded one ethanol control frog from analysis. Using R version 4.5.1 (R Core Team, 2025), we compared the number of mouth-gape counts per animal before and after solution administration within each treatment using a Wilcoxon test (package ‘stats’ version 4.5) due to non-normal distribution of residuals. We also compared the number of mouth-gape events after solution administration among the treatments using a Kruskall-Wallis test (package ‘rstatix’ version 0.7, Kassambara, 2023). We created the graphs using packages ‘ggplot2’ version 4.0.0 (Wickham, 2016), ‘ggbeeswarm’ version 0.7.3 (Clarke et al., 2025), ‘ggpubr’ version 0.6.1 (Kassambara, 2025), and ‘RColorBrewer’ version 1.1-3 (Neuwirth, 2022).

### 2.4 Whole-animal performance assays

We evaluated locomotion and behavior using an arena similar to assays previously used to evaluate resistance to toxins in snakes and lizards (Brodie and Brodie, 1990; Ridenhour et al., 2004; Thill et al., 2025). This bioassay assumes that resistant animals will maintain baseline performance after exposure to a specific dose of toxins, while non-resistant animals will decrease their performance. In our assays, we measured several locomotion and behavioral parameters that could be affected by disrupted muscular and neuronal nAChR function (Badio and Daly, 1994; Tarvin et al., 2017a). We evaluated how long it took them to react after being placed in the arena, how long they rested between jumps, how far they jumped, and how active they were in the arena. We used a repeated measures design to measure the performance of each individual at four time points after solution administration: 3 minutes, 1 hour, 4 hours, and 24 hours. We then compared the performance of individuals exposed to epibatidine to the baseline behavior of animals in the ethanol and water control groups.

We placed the racetrack (45.7 cm x 12.7 cm, Fig. 1B) inside a plastic container of 40 cm height, and a smart phone (iPhone 11 [iOS 18.1.1], iPhone 12 [iOS 26.3], or iPhone 13 [iOS 26.3]) on a panel on top of the box to record the arena. All recordings were taken at 30 frames per second (fps). For the measurement of 3 minutes after solution administration, we kept the individuals for 2 minutes in individual transparent cups (7.5 cm diameter) for close observation of distress behavior and mouth-gaping recording. We then turned the cup upside down onto the starting area of the arena and covered it with an opaque cup to allow the frog to acclimate. After an additional minute, we started the recording, lifted the cup, and allowed the frog to move freely in the arena. We stopped the recording as soon as the frog crossed the opposite line of the arena. If a frog did not move after approximately 1 minute, we used a flat piece of cardboard and moved it 1.5 cm each 10 seconds to stimulate the frog. After the 3-minute trial, we kept the frogs in individual cups in a dark environment to limit additional stress. We repeated the process of turning the cup upside down, covering it with an opaque cup, and waiting for 1 minute before starting the 1-hour time point. After the 1-hour trial, we put the frogs back in their terraria. Shortly before the 4-hour and 24-hour timepoints, we collected animals from their terraria, identified them, and repeated the protocol (Fig. 1C).

### 2.5 Video analysis

All videos were converted to an 8-bit, greyscale image stack using FFmpeg version 6.0 (Tomar, 2006). Individual image stacks were then uploaded into Fiji version 2.17.0 for analysis (Schindelin et al., 2012). For each stack, we set the scale using one centimeter markings drawn on the racetrack. Using the Manual Tracking plugin, we quantified the frog’s path through the racetrack by clicking on the tip of the animal’s nose in each frame. We began tracking on the first frame where the frog’s nose was visible, and ended when the frog had entirely crossed the end of the racetrack. In the event that a frog jumped out of frame or its nose was not visible, we clicked at the last visible point until the frog reappeared. In frames where the nose was blurry, such as during a quick jump, we clicked on the middle of the blurred area created by the nose. We noted if the frog jumped out of frame or if cardboard was needed to stimulate the frog to jump. When the track was completed, we saved the output as a CSV file containing the frog’s coordinates, distance traveled, and velocity for each frame. If we suspected that the frog’s behavior was compromised due to any external factor other than the solution administration (*e.g.*, frog escaped from the arena and needed to be recaptured, affecting its baseline behavior, or frog went out of frame mid crossing, impacting our ability to measure jump distance or frequency), we removed the trial from further analysis. Additionally, due to technical issues, we did not have recordings of 13 trials (detailed list of videos included in the analyses is available in Supplemental Material 2).

We analyzed video data with a custom R script using the packages ‘tidyverse’ version 2.0.0 (Wickham et al., 2019) and ‘janitor’ version 2.2.1 (Firke et al. 2024). For each CSV file, we calculated four parameters. First, the latency to first reaction was calculated as the duration in seconds from the initial frame to the first frame where velocity exceeded the reaction threshold of > 9.0 cm/s. We then annotated individual jumping events as a block of frames with continuous movement > 5 cm/s and peak velocity exceeding 20 cm/s. Using these data we calculated the following additional parameters: number of jumps per second, by dividing the total number of jumps by the total time spent in the arena from the moment the cup was lifted until they crossed the finish line (*i.e.*, number of frames up to the finish line divided by the frame rate of 30 fps), average distance per jump (centimeters), and average time between jumping events (seconds).

To assess the impact of epibatidine on behavior and locomotion, we compared metrics of frog behavior between the ethanol control and the two epibatidine treatments across all four time points (3 min, 1 h, 4 h, and 24 h). To analyze latency to first reaction, jumps per second, distance of jumps, and time between jumps, we used a generalized linear mixed model in R to predict each metric (GLMM, *glmmTMB* function from package ‘glmmTMB’ version 1.1.14, Brooks et al., 2017) with a gamma distribution and a log-link function to account for the right-skewed nature of the data. Each model included treatment, time point, and their interaction as fixed effects, stimulation (the need of the flat piece of cardboard to stimulate movement) and frog mass as covariates, and individual as a nested random effect. We also tested the impact of ethanol and ethanol+epibatidine by including the water control samples. We use the same model set-up described above but excluded the 24-hour trials because of low sample size for the water control (N = 2).

To test for the impact of alkaloid accumulation on animal locomotion and behavior, we used the same model described above, with epibatidine treatment groups only (2.0 and 6.0 µg, N = 5 per epibatidine treatment). We also included epibatidine quantity in ng/mg of wet skin mass as a covariate.

For all GLMM analyses, we checked model assumptions using residual diagnostics in the package ‘DHARMa’ (Hartig, 2024). In some cases, the residuals deviated from the expected distribution due to differential residual dispersion of frog mass. In these cases, we fixed the model by adjusting the variance of the frog mass using the ‘*dispformula*’ argument in *glmmTMB*. In a few cases, where adjusting the variance was not enough to fit the model in the necessary assumptions, we log transformed the data and used a Gaussian distribution (rather than gamma with log-link function) in the model. The overall significance of the fixed effects and covariates were tested using type II Wald chi-square tests (‘car’ package version 3.1-3, (Fox and Weisberg, 2019). When necessary, we ran pairwise Tukey HSD post-hoc comparisons using the ‘emmeans’ package version 1.11.2-8 (Lenth, 2025). Whenever the stimulation with the flat piece of cardboard had a significant effect on the focal parameter (except for latency to first reaction; see Results), we removed these frogs from the analysis, and reran the model. We created the graphs using the same R packages as in section 2.3.

### 2.6 Epibatidine quantification

Approximately 30 minutes after the 24-hour trial, we euthanized a subset of the frogs (5 samples of each epibatidine treatment) by cooling the frogs for 10 minutes in the fridge and flash freezing in liquid nitrogen (Lillywhite et al., 2017; Shine et al., 2015). We collected half of the dorsal and ventral skin, measured their mass (Mettler Toledo MA104, ± 0.1 mg), and stored them individually in glass vials with Teflon-lined caps containing 500 µL of 100% methanol. We collected 100 µL of each skin methanol extract into 1.5 mL microcentrifuge tubes, centrifuged them at 13,000 rpm at 4°C for 15 minutes, and collected the supernatant in 2-mL GC-MS autosampler vials with an insert. Then, we added 100 ng of nicotine ((−)-nicotine ≥ 99%, CAS: 54-11-5, Sigma-Aldrich, Darmstadt, Germany) as an internal standard, evaporated the methanol to dryness under nitrogen flow, and resuspended the alkaloids with 10 µL of 100% methanol for gas chromatography-mass spectrometry (GC-MS) analysis.

We performed GC-MS on an Agilent GC 7890A and MS 5975C. The GC was fitted with a fused silica capillary column (Agilent, DB-5MS, 30 m × 0.32 mm × 0.25 µm) with helium as the carrier gas (1.28 mL/min). Extracts were analyzed in pulsed splitless mode, with a 1 µL injection volume, and a column oven temperature program of 100°C for 4 min, then increased by 10°C min−1 to 250°C. Injector and transfer line temperatures were maintained at 280°C. To yield cleaner profiles, we set up a Selected Ion Monitoring (SIM) method, narrowing the mass range to the nicotine (m/z: 84, 133, 162, 161, and 42) and epibatidine (m/z: 69, 68, and 140) specific ions. We identified both compounds by comparing MS properties and GC retention times with standard solutions run using the same method and instrument. We determined the quantity of epibatidine by comparing the observed alkaloid peak areas to the peak area of the nicotine internal standard, using OpenChrom 1.5.0 (Wenig and Odermatt, 2010). We detected chromatogram peaks using the Peak Detector feature and integrated each peak area using the Trapezoid Peak Integrator feature.

We summed the total amount of alkaloid detected in dorsal and ventral skin and multiplied it by two as we quantified only half of the skin and calculated two metrics from the alkaloid quantification. First, we divided the total quantity of epibatidine by the wet skin mass to measure epibatidine accumulation capacity (ng/mg), and then we calculated the percentage of epibatidine detected in the skin from the total amount fed to measure the efficiency of accumulation. We then compared both metrics between the 2-µg and 6-µg treatments using the Wilcoxon test (package ‘stats’ version 4.5, R Core Team, 2025). We created the graphs using the same methods as section 3.2.

## 3. RESULTS

None of the frogs presented the mouth-gaping behavior before solution administration, but after administration, several frogs presented mouth-gaping behavior (Fig. 2A). When comparing within treatments, only the 6-µg treatment showed a significant increase in mouth-gaping behavior after the alkaloid solution administration (p = 0.0035, Fig. 2B). However, when comparing the mouth-gaping behavior after administration among all treatments, there was no significant difference (p = 0.47).

**Figure 2.**
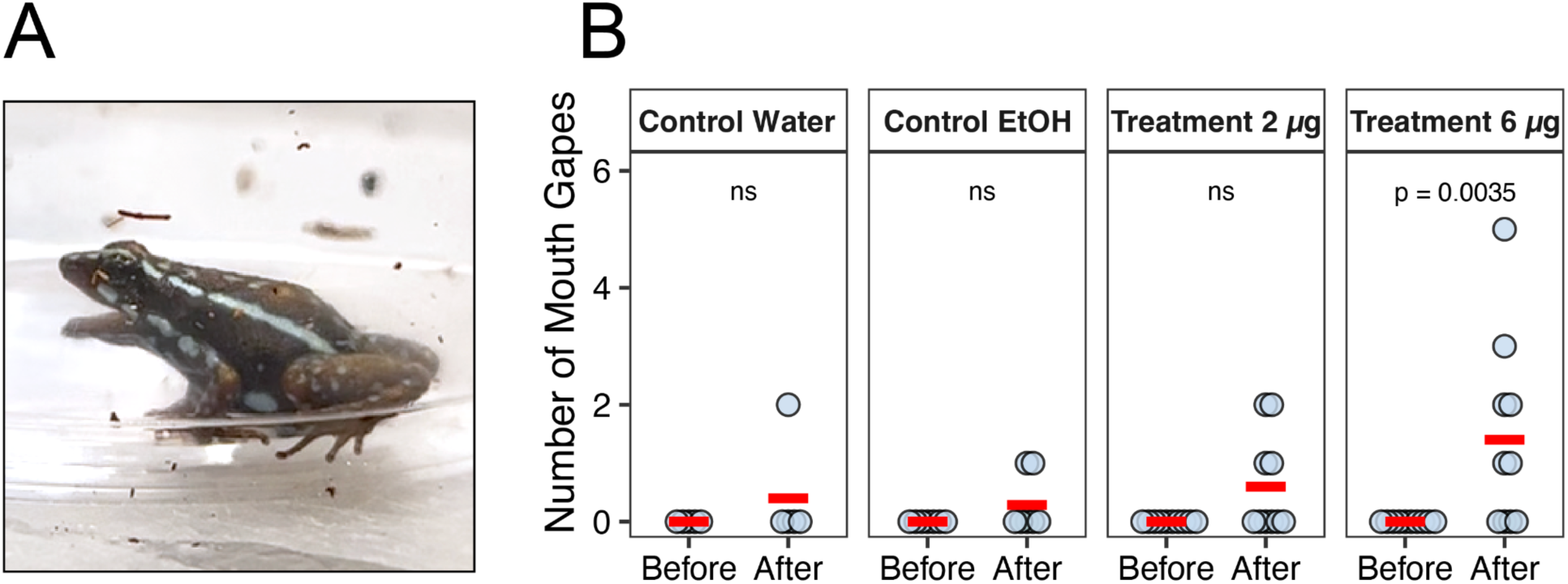
Mouth-gaping behavior 20 seconds before and 20 seconds after solution administration. A) *Epipedobates tricolor* performing mouth-gaping behavior. B) Number of mouth gapes by each animal in each treatment. There was no difference among all “after” mouth gape counts (Kruskall-Wallis, p = 0.47). Each circle represents an individual frog and the red trace represents the mean number for each group. ns = not significant.

During the whole-animal performance assays, *E. tricolor* frogs readily traversed the arena. We removed six trials from the analyses (one 6-µg treatment at 4 h, one 6-µg treatment at 24 h, two ethanol control at 1 h, one ethanol control at 4 h, and one water control at 3 min) due to behavior compromised by external factors (e.g., frogs escaping from or jumping outside of the arena). In the analysis that included all experimental groups, the water control group tended to have lower average latency to first reaction and longer average distance per jump at all time points. However, these differences were statistically significant only at the 4-hour time point (Fig. 1A–B, Table SM1–3 in Supplemental Material 1).

In the analysis testing for the impact of epibatidine on behavior, 15 out of the 114 trials needed cardboard to stimulate movement, which included 10 different individuals (Fig. 3E). The need for external stimulation in these frogs significantly affected the latency to first reaction, number of jumps for second, and the average time between jumps (Table 1). Due to the nature of latency to first reaction parameter and to avoid biasing the dataset toward the most active individuals, no frogs were removed from this analysis; for the other two parameters (number of jumps per second and average time between jumps) the frogs that needed stimulation to move were removed from analyses. Ultimately, there were no significant effects of treatment, time point, their interaction, or body mass on any of the parameters tested (Table 1). However, body mass significantly influenced the variance of first reaction time (p = 0.002), with larger individuals exhibiting significantly higher behavioral variability, demonstrating that body mass directly affects the behavioral predictability in this case.

**Figure 3.**
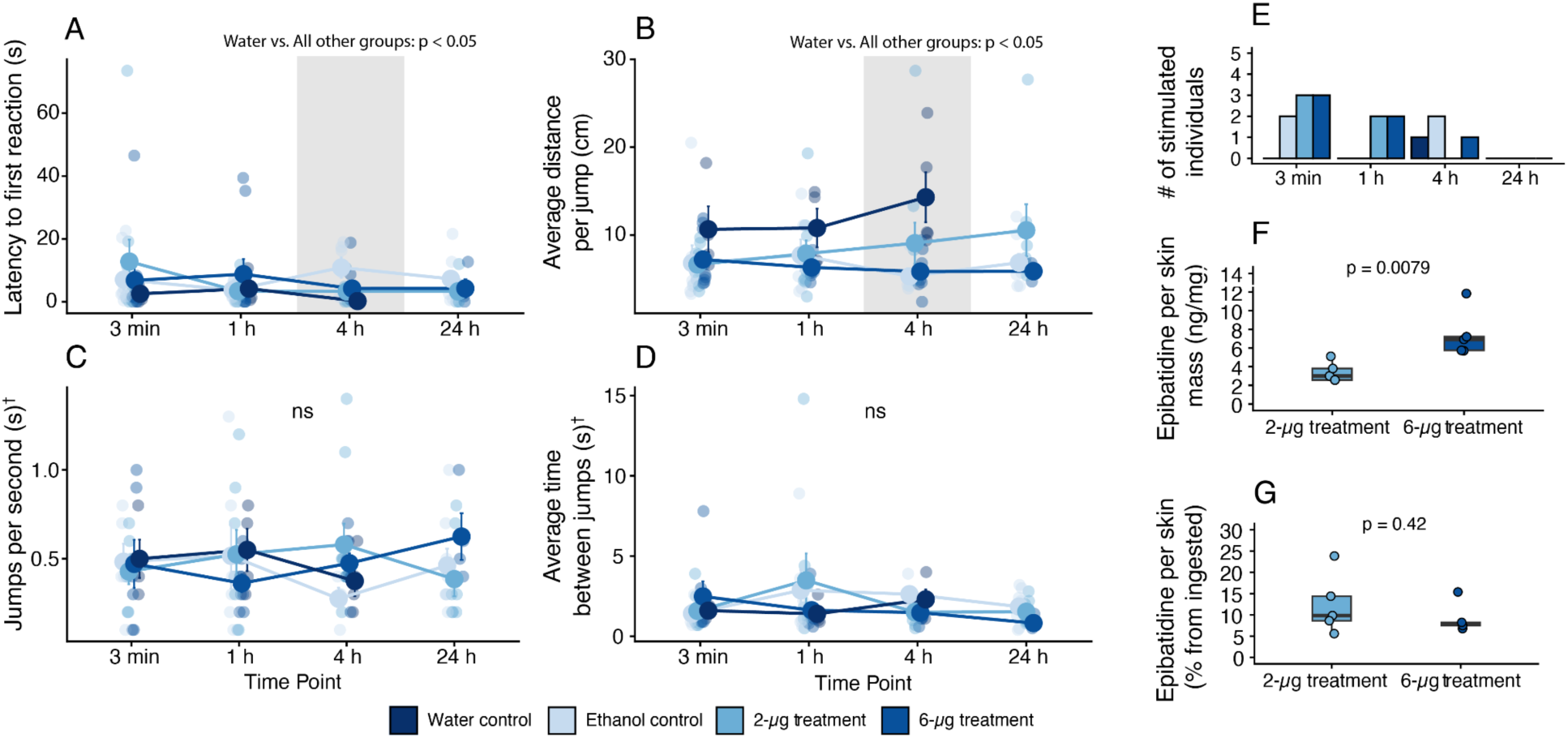
Impact of different doses of epibatidine on reaction and locomotion at different times points after solution administration. A) Latency for the frog to make a first reaction; B) Average distance in centimeters per jump; C) Number of jumps per second; D) Average time in seconds between jumps; E) Number of individuals that needed stimulation to move after 60 seconds from the start of the experiment; F) Quantity of epibatidine (ng) per skin mass (mg) of individuals from 2-µg treatment and 6-µg treatment; and G) Percentage of epibatidine detected in the skin compared to the total quantity administered. Treatments, Time Points, or the interaction between them did not have effect in any of the parameters when comparing only ethanol control vs. epibatidine controls (GLMM, p > 0.05). When water control is added to the model, there is a significant difference in latency for first reaction and average distance per jump in the 4-hour time point between water and all other groups (grey shaded areas). †: Stimulated frogs removed from analysis.

**Table 1.** Results from the generalized linear mixed models including ethanol control and epibatidine treatment samples. Latency to first reaction and average distance per jump includes stimulated frog trials, whereas jumps per second and average time between jumps do not due to significant effects of stimulation in the models. See Table SM1 in Supplemental Material 1 for models including water controls.

| Parameter | Coefficient | X<sup>2</sup> | d.f. | p value |
| --- | --- | --- | --- | --- |
| Latency to first reaction | Treatment | 0.557 | 2 | 0.757 |
|  | Time | 0.799 | 3 | 0.850 |
|  | Body mass | 0.031 | 1 | 0.860 |
|  | Stimulation | 4.976 | 1 | **0.026** |
|  | Treatment:Time | 2.626 | 6 | 0.854 |
| Average distance per jump (cm) | Treatment | 2.235 | 2 | 0.327 |
|  | Time | 2.315 | 3 | 0.510 |
|  | Body mass | 0.090 | 1 | 0.765 |
|  | Stimulation | 0.186 | 1 | 0.666 |
|  | Treatment:Time | 8.694 | 6 | 0.192 |
| Jumps per second | Treatment | 0.339 | 2 | 0.844 |
|  | Time | 0.041 | 3 | 0.998 |
|  | Body mass | 0.001 | 1 | 0.979 |
|  | Stimulation | Stimulated frogs removed from analysis |  |  |
|  | Treatment:Time | 8.630 | 6 | 0.196 |
| Average time between jumps | Treatment | 2.349 | 2 | 0.309 |
|  | Time | 3.247 | 3 | 0.355 |
|  | Body mass | 1.398 | 1 | 0.237 |
|  | Stimulation | Stimulated frogs removed from analysis |  |  |
|  | Treatment:Time | 5.469 | 6 | 0.485 |

Quantification of epibatidine in the skin showed a dose-dependent trend, with the 2-µg treatment frogs having significantly less epibatidine per milligram of skin than the 6-µg treatment frogs (2-µg treatment = 3.4 ± 1.1 ng/mg, 6-µg treatment = 7.5 ± 2.5 ng/mg, p = 0.0079, Fig. 3F). However, there was no significant difference when calculating the percentage of epibatidine detected relative to the total amount fed (2-µg treatment = 13.0% ± 6.5%, 6-µg treatment = 9.1% ± 3.5%, p = 0.42, Fig 3G). We found no effect of alkaloid quantity per skin mass on any of the parameters analyzed in the subset of frogs for which epibatidine was quantified (Table SM4 in Supplemental Material 1).

## 4. DISCUSSION

Our results demonstrate that *Epipedobates tricolor* maintains locomotor performance and behavior following exposure to different doses of epibatidine. The analyzed parameters are representative of physiological processes, such as respiration and muscular activity, which could be impacted by epibatidine’s binding activity to nAChR (Salehi et al., 2018). The absence of differences between epibatidine and ethanol control treatments highlights these frogs’ resistance capacity, even when consuming high quantities of epibatidine. This suggests that this species avoids a physiological trade-off between toxin ingestion and the defensive benefits of sequestration.

Mouth-gaping is a recognized stress indicator in animals (*e.g.*, Taylor et al., 2021). Observing this behavior only after solution administration, along with the lack of significant differences in mouth-gaping behavior among treatments, indicates that the main stressors in this study are handling and solution administration, not epibatidine. However, there is an increasing trend in the number of individuals presenting mouth-gaping behavior from the water control to the 6-µg treatment (Fig. 2B). Only one of five animals given water control was observed mouth gaping, while two of eight frogs in the ethanol control group, four of ten frogs from 2-µg treatment group, and six of ten frogs from 6-µg treatment mouth gaped after solution administration. Mouth gaping may indicate buccal respiration, which is unusual for frogs (Taylor et al., 2010), or a response to the unpalatability of the compound (Hurlbert, 1970). Amphibians possess a variety of bitter taste receptors, including on the tongue, and lipophilic compounds, such as epibatidine and other alkaloids, are perceived as bitter (Higgins et al., 2025). Ethanol may also trigger a reaction due to its unpleasant burning sensation. Mouth gaping following ingestion of distasteful or toxic substances has been documented in various species, including frogs, snakes, and birds (Hurlbert, 1970).

A higher number of individuals needed stimulation to start moving at the 3-minute time point than at all other time points, especially the 24-hour time point (Fig. 3E). This may reflect stress from handling and oral administration (Ricciardella et al., 2010), or changes in behavior due to gradual acclimation to the arena environment (Barnett et al., 2025; Blanchette et al., 2017; Soto et al., 2024). For instance, studies on other dendrobatid species have shown that latency to explore a new environment decreases with repeated trials (Soto et al., 2024). Additionally, a frog’s personality and propensity for movement may influence this behavior (Soto et al., 2024), as four of ten individuals requiring stimulation needed it at multiple time points.

The absence of negative effects of high epibatidine doses on the locomotor capacity and behavior of *E. tricolor in vivo* is consistent with the *in vitro* molecular resistance reported in previous studies (Tarvin et al., 2017a; York et al., 2023). It remains unclear whether other nAChR isoforms possess substitutions that contribute to resistance against high levels of epibatidine. In vertebrates, neuronal isoforms typically consist of pentameric α4β2 combinations, where the ACh binding site is located at the αβ interface. The substitutions in *Epipedobates* spp. are located in the β-subunit of this neuronal isoform (Tarvin et al., 2017a). Conversely, the adult muscle isoform consists of α2βεγ subunits, with ACh binding at the αγ and αε interfaces (Hille, 2001; Kalamida et al., 2007). It is unclear whether mutations in the β-subunit of the muscle isoform would confer equivalent resistance given the structural differences in the binding sites of the neuronal and muscle isoforms. The lack of significant performance costs suggests either that the current resistance mechanisms are highly effective or that yet-to-be-described substitutions in other nAChR subunits allow for full resistance, resulting in an absence of measurable physiological costs.

In contrast to our findings, previous studies have reported physiological costs associated with higher doses of epibatidine in *Epipedobates anthonyi*. Waters et al., (2023) observed lethargy and immobility for about 30 minutes after a 10-µg dose in a 50% ethanol vehicle was administered to a 0.4-g *E. anthonyi*. Our maximum dose was lower (6 µg), and the larger body mass of our *E. tricolor* (0.5–0.8 g) likely explains why we did not observe the same extreme physiological response. Additionally, our alkaloid solutions contained 2–6% ethanol compared to the 50% ethanol solutions used in their study and in previous alkaloid administration studies (*e.g.*, Jeckel et al., 2026, 2022; Minder et al., 2026). We found differences in latency to the first reaction and jump distance in the 4-hour time point between the water control group and all the other groups, in which ethanol was present. This shows that even small percentages of ethanol, such as 2% and 6%, may significantly impact physiological processes in poison frogs. Further studies are needed to understand the effects of ethanol on the physiological processes and sequestration mechanisms of poison frogs, as well as to establish alternative methods of administering lipophilic compounds, such as epibatidine. Notably, the *E. anthonyi* in the Waters et al. (2023) study survived and were able to continue the experiment at a lower dose (5 µg). This further supports a degree of resistance to epibatidine and/or ethanol.

Other forms of resistance may play an important role in the ability to ingest toxins (Ramírez-Castañeda et al., 2025). The low percentage of epibatidine detected in the skin (∼10%) is consistent with previous alkaloid feeding studies, which indicate that only some of the ingested alkaloids remain unmodified in the body (Jeckel et al., 2026, 2022; Waters et al., 2023). Metabolism and detoxification of ingested toxins are common resistance strategies that are present even in sequestering organisms (Heckel, 2014).

One such example is the pyrrolizidine alkaloid detoxification mechanism in arctiid moths. They *N*-oxidize the toxic reduced tertiary form of the alkaloid back to its stable non-toxic hydrophilic form via a flavin-dependent monooxygenase enzyme pathway, and these *N*-oxidized alkaloids are safely stored in tissues throughout the pupal and adult stages (Hartmann and Ober, 2000; Naumann et al., 2002). In poison frogs, several CYP450 proteins and *N*-methylation enzymes are down-regulated in alkaloid-fed frogs, while some other CYP450s and a glutathione *S*-transferase are up-regulated (O’Connell et al., 2021). Although the metabolism of alkaloids in poison frogs that sequester alkaloids is not well understood, these examples combined with feeding trial data (this study, Waters et al., 2023) demonstrate that there must be a balance between maintaining intact alkaloids for defense and alleviating the physiological burden of high toxin load.

In wild *Epipedobates* and *Ameerega* frogs, epibatidine is present in much lower concentrations than other alkaloids. The methanol extract from the skins of 750 *E. anthonyi* individuals collected near Santa Isabel, Azuay, Ecuador in 1976 contained approximately 21 mg of pumiliotoxin **251D** and only about 1 mg of epibatidine (Daly et al., 2000; Jain et al., 1995). Similarly, epibatidine levels in *E. anthonyi* from Moromoro, El Oro, Ecuador were 37 times lower than those of 5,8-indolizidine **231C**, a prevalent alkaloid in several poison frogs (Daly et al., 2005; Tarvin et al., 2024). Since the specific arthropod source of epibatidine is unknown, it is unclear how much metabolism and availability interfere with epibatidine sequestration. However, the potency of epibatidine implies that even small quantities can have adverse effects on predators and could provide sufficient chemical protection (Badio and Daly, 1994).

Additionally, when discussing toxicity of defensive compounds, the method of administration must be considered as it directly impacts the exposure of target receptors to the toxin. With oral administration, enzymes in the digestive tract and liver may metabolize some of the toxin. This explains the high toxicity and bioavailability observed with intravenous or subcutaneous exposure, which evades digestive and detoxification enzyme activity (Badio and Daly, 1994; Williams et al., 2002). However, oral administration of TTX in *T. sirtalis* snakes resulted in a longer toxin half-life compared to subcutaneous injection (Robinson et al., 2024). This could be explained by the difference in diffusion time from the food to the digestive tract (Robinson et al., 2024), or an adaptation to retain TTX for defensive purposes in the snakes (REF). In the case of epibatidine, the intravenous lethal dose (LD₅₀) in mice is approximately 2 µg/kg (Badio and Daly, 1994), suggesting that the frogs in our study are nearly 5,000 times more resistant (6 µg/0.6 g). However, our frogs received epibatidine orally with a 2–6% ethanol vehicle, so it likely underwent some form of metabolism. Remarkably, *E. tricolor* is still 20 times more resistant than the oral LD₅₀ of 5.1 mg/kg provided by the epibatidine manufacturer (test animal unknown; Sigma-Aldrich, Darmstadt, Germany).

Further evidence of the likelihood of the presence of additional epibatidine resistance strategies in *E. tricolor*, such as metabolism, comes from other dendrobatid species lacking TSR, including some *Ranitomeya*, *Phyllobates*, and *Dendrobates* species (Tarvin et al., 2017a), which were able to ingest 10 µg of epibatidine for seven consecutive days (*i.e.*, 70 µg in total), and sequester 9–20% of it in their skin (Waters et al., 2023). Although the study did not explicitly test for physiological costs, the results demonstrate TSR is not strictly necessary for surviving and sequestering epibatidine.

Other resistance mechanisms in poison frogs may include specialized proteins such as saxiphilin, a STX-binding protein (Abderemane-Ali et al., 2021), and alkaloid-binding globulins (ABG; Alvarez-Buylla et al., 2023). These proteins readily bind alkaloids and may function as plasma transporters and alternative targets. Interestingly, ABG did not bind to epibatidine in *Epipedobates tricolor*, highlighting the diversity of mechanisms these frogs have to resist and/or sequester specific alkaloids.

## 5. CONCLUSION

Our findings demonstrate that *E. tricolor* exhibits significant resistance to orally administered epibatidine. Even at doses much higher than those expected in nature, the frog experiences minimal effects on locomotor performance and behavior. These results support the hypothesis that poison frogs have mechanisms that reduce the physiological consequences of alkaloid sequestration. Additionally, our results suggest that the observed behavioral shifts, such as mouth-gaping and the need for stimulation to move, are likely primarily driven by handling stress and sensitivity to the ethanol solution rather than epibatidine. These results highlight the need for further dendrobatid research to establish a non-alcoholic delivery vehicle.

Furthermore, the low recovery of epibatidine from the skin and the considerable difference between the observed resistance in *E. tricolor* and baseline mammalian toxicity models indicates multiple physiological and molecular mechanisms of resistance and sequestration. These mechanisms may include TSR, metabolism, and alternative target molecules, among others. Finally, we have demonstrated that *E. tricolor* is resistant to epibatidine *in vivo*, which is consistent with its previously demonstrated resistance *in vitro*. Like other animals that specialize in toxic food, their ability to ingest large quantities of toxins indicates an adaptive, multi-faceted response that enables resistance, sequestration, and consequently, chemical defense.

## Acknowledgments

Alyssa Bui, Chenhey Chu, Indra Deshmukh, Camille Dayton, Evan Ho, and Alina Tran assisted with frog husbandry. Ziting Chen assisted with oral administration. Alyssa Bui assisted with sample preparation for GC-MS. Ziting Chen, Ammon Corl, María José Navarrete, Kannon Pearson provided valuable suggestions in the first drafts of the manuscript.

## Funding

This study was supported by NIH NIGMS R35GM150574 to RDT, which supported salaries of RDT, SJD, and AMJ, as well as research expenses. GCMS work, NDT, and RBW were supported by USDA Hatch Project (CA-B-INS-0335-H) to NDT. Behavioral assays were supported by the Integrative Biology Summer Undergraduate Research Experience (SURE) program from the Department of Integrative Biology, University of California, Berkeley.

## Data availability

All videos, raw data and analysis code are available in the Dryad data repository at https://doi.org/10.5061/dryad.41ns1rnwm

## AI statement

The R script for analyzing video data was written in collaboration with Gemini v3.1 Pro.

## Notes

https://doi.org/10.5061/dryad.41ns1rnwm

### Competing Interest Statement

The authors have declared no competing interest.

## References

Abderemane-Ali, F., Rossen, N.D., Kobiela, M.E., Craig, R.A., II, Garrison, C.E., Chen, Z., Colleran, C.M., O’Connell, L.A., Du Bois, J., Dumbacher, J.P., Minor, D.L., Jr., 2021. Evidence that toxin resistance in poison birds and frogs is not rooted in sodium channel mutations and may rely on “toxin sponge” proteins. J. Gen. Physiol. 153, e202112872. 10.1085/jgp.202112872

Agrawal, A.A., Böröczky, K., Haribal, M., Hastings, A.P., White, R.A., Jiang, R.-W., Duplais, C., 2021. Cardenolides, toxicity, and the costs of sequestration in the coevolutionary interaction between monarchs and milkweeds. Proc. Natl. Acad. Sci. 118, e2024463118. 10.1073/pnas.2024463118

Alvarez-Buylla, A., Fischer, M.-T., Moya Garzon, M.D., Rangel, A.E., Tapia, E.E., Tanzo, J.T., Soh, H.T., Coloma, L.A., Long, J.Z., O’Connell, L.A., 2023. Binding and sequestration of poison frog alkaloids by a plasma globulin. eLife 12, e85096. 10.7554/eLife.85096

Badio, B., Daly, J.W., 1994. Epibatidine, a potent analgetic and nicotinic agonist. Mol. Pharmacol. 45, 563–569.

Barnett, J.B., McEwen, B.L., Kinley, I., Anderson, H.M., Yeager, J., 2025. Behavioural mimicry among poison frogs diverges during close-range encounters with predators. J. Evol. Biol. 38, 663–671. 10.1093/jeb/voaf038

Blanchette, A., Becza, N., Saporito, R.A., 2017. Escape behaviour of aposematic (*Oophaga pumilio*) and cryptic (*Craugastor sp*.) frogs in response to simulated predator approach. J. Trop. Ecol. 33, 165–169. 10.1017/S0266467417000037

Brodie, E.D., III, Brodie, E.D., Jr., 1990. Tetrodotoxin resistance in garter snakes: an evolutionary response of predators to dangerous prey. Evolution 44, 651–659. 10.1111/j.1558-5646.1990.tb05945.x

Brodie III, E.D., Brodie Jr., E.D., 1999. Costs of exploiting poisonous prey: evolutionary trade-offs in a predator-prey arms race. Evolution 53, 626–631. 10.1111/j.1558-5646.1999.tb03798.x

Brooks, M., Kristensen, K., Benthem, K. van, Magnusson, A., Berg, C., Nielsen, A., Skaug, H., Mächler, M., Bolker, B., 2017. glmmTMB balances speed and flexibility among packages for zero-inflated generalized linear mixed modeling. R J.

Brower, L.P., Ryerson, W.N., Coppinger, L.L., Glazier, S.C., 1968. Ecological chemistry and the palatability spectrum. Science 161, 1349–1350.

Camara, M.D., 1997. Physiological mechanisms underlying the costs of chemical defence in *Junonia coenia* Hubner (Nymphalidae): A gravimetric and quantitative genetic analysis. Evol. Ecol. 11, 451–469. 10.1023/A:1018436908073

Caty, S.N., Alvarez-Buylla, A., Byrd, G.D., Vidoudez, C., Roland, A.B., Tapia, E.E., Budnik, B., Trauger, S.A., Coloma, L.A., O’Connell, L.A., 2019. Molecular physiology of chemical defenses in a poison frog. J. Exp. Biol. 222. 10.1242/jeb.204149

Clarke, E., Sherrill-Mix, S., Dawson, C., 2025. ggbeeswarm: categorical scatter (violin point) plots. 10.32614/CRAN.package.ggbeeswarm

Daly, J.W., Garraffo, H.M., Spande, T.F., Decker, M.W., Sullivan, J.P., Williams, M., 2000. Alkaloids from frog skin: the discovery of epibatidine and the potential for developing novel non-opioid analgesics. Nat. Prod. Rep. 17, 131–135. 10.1039/a900728h

Daly, J.W., Martin Garraffo, H., Spande, T.F., Jaramillo, C., Stanley Rand, A., 1994a. Dietary source for skin alkaloids of poison frogs (Dendrobatidae)? J. Chem. Ecol. 20, 943–955. 10.1007/BF02059589

Daly, J.W., Secunda, S., Garraffo, H.M., Spande, T.F., Wisnieski, A., Cover Jr, J.F., 1994b. An uptake system for dietary alkaloids in poison frogs (Dendrobatidae). Toxicon 32, 657–663.

Daly, J.W., Spande, T.F., Garraffo, H.M., 2005. Alkaloids from amphibian skin: a tabulation of over eight-hundred compounds. J. Nat. Prod. 68, 1556–1575. 10.1021/np0580560

Douglas, T.E., Beskid, S.G., Gernand, C.E., Nirtaut, B.E., Tamsil, K.E., Fitch, R.W., Tarvin, R.D., 2022. Trade-offs between cost of ingestion and rate of intake drive defensive toxin use. Biol. Lett. 18, 20210579. 10.1098/rsbl.2021.0579

Dussourd, D.E., 1999. Behavioral sabotage of plant defense: do vein cuts and trenches reduce insect exposure to exudate? J. Insect Behav. 12, 501–515. 10.1023/A:1020966807633

Dussourd, D.E., Eisner, T., 1987. Vein-cutting behavior: insect counterploy to the latex defense of plants. Science 237, 898–901. 10.1126/science.3616620

Feldman, C.R., Brodie Jr, E.D., Brodie III, E.D., Pfrender, M.E., 2012. Constraint shapes convergence in tetrodotoxin-resistant sodium channels of snakes. PNAS 109, 4556–4561. 10.1073/pnas.1113468109/-/DCSupplemental.www.pnas.org/cgi/doi/10.1073/pnas.1113468109

Fox, J., Weisberg, S., 2019. An R companion to applied rgression, Third. ed. Sage, Thousand Oaks CA.

Geffeney, S., Brodie, E.D., Ruben, P.C., Brodie, E.D., 2002. Mechanisms of adaptation in a predator-prey arms race: TTX-resistant sodium channels. Science 297, 1336–1339. 10.1126/science.1074310

Geffeney, S.L., Fujimoto, E., Brodie Jr, E.D., Brodie III, E.D., Ruben, P.C., 2005. Evolutionary diversitication of TTX-resistant sodium channels in a predator-prey interaction. Nature 434, 759–63. 10.1038/nature03392

Goetz, S.M., Guyer, C., Boback, S.M., Romagosa, C.M., 2018. Toxic, invasive treefrog creates evolutionary trap for native gartersnakes. Biol. Invasions 20, 519–531. 10.1007/s10530-017-1554-6

Hague, M.T.J., Toledo, G., Geffeney, S.L., Hanifin, C.T., Brodie, E.D., Jr., Brodie, E.D., III, 2018. Large-effect mutations generate trade-off between predatory and locomotor ability during arms race coevolution with deadly prey. Evol. Lett. 2, 406–416. 10.1002/evl3.76

Hartig, F., 2024. DHARMa: Residual diagnostics for hierarchical (multi-level / mixed) regression models. 10.32614/CRAN.package.DHARMa

Hartmann, T., Ober, D., 2000. Biosynthesis and metabolism of pyrrolizidine alkaloids in plants and specialized insect herbivores. Top. Curr. Chem. 209, 208–243. 10.1007/3-540-48146-X

Heckel, D.G., 2014. Insect detoxification and sequestration strategies, Annual Plant Reviews online. 10.1002/9781119312994.apr0507

Higgins, K.W., Itoigawa, A., Toda, Y., Bellott, D.W., Anderson, R., Márquez, R., Weng, J.-K., 2025. Rapid expansion and specialization of the TAS2R bitter taste receptor family in amphibians. PLOS Genet. 21, e1011533. 10.1371/journal.pgen.1011533

Hille, B., 2001. Ionic channels of excitable membranes, 3rd ed. Sinauer Associates Inc., Sunderland.

Hurlbert, S.H., 1970. Predator responses to the Vermilion-Spotted Newt(*Notophthalmus viridescens*). J. Herpetol. 4, 47–55. 10.2307/1562702

Jain, P., Garraffo, H.M., Spande, T.F., Yeh, H.J.C., Daly, J.W., 1995. A new subclass of alkaloids from a dendrobatid poison frog: a structure for deoxypumiliotoxin 251H. J. Nat. Prod. 58, 100–104. 10.1021/np50115a012

Jeckel, A.M., Bolton, S.K., Waters, K.R., Antoniazzi, M.M., Jared, C., Matsumura, K., Nishikawa, K., Morimoto, Y., Grant, T., Saporito, R.A., 2022. Dose-dependent alkaloid sequestration and N-methylation of decahydroquinoline in poison frogs. Journal of Experimental Zoology Part A: Ecological and Integrative Physiology 1–10. 10.1002/jez.2587

Jeckel, A.M., Saporito, S.K., Antoniazzi, M.M., Jared, C., Matsumura, K., Nishikawa, K., Morimoto, Y., Grant, T., Saporito, R.A., 2026. Experimental evidence supports gradual evolution of alkaloid sequestration in poison frogs. Proc. R. Soc. B Biol. Sci. 293, 20253144. 10.1098/rspb.2025.3144

Jones, T.H., Blum, M.S., 1982. Ant venom alkaloids from *Solenopsis* and *Monomorium* species. Tetrahedron 38, 1949–1958. 10.1016/0040-4020(82)80044-6

Jones, T.H., Torres, J.A., Spande, T.F., Garraffo, H.M., Blum, M.S., Snelling, R.R., 1996. Chemistry of venom alkaloids in some *Solenopsis* (Diplorhoptrum) species from Puerto Rico. J. Chem. Ecol. 22, 1221–1236. doi:10.1007/BF02266962

Kalamida, D., Poulas, K., Avramopoulou, V., Fostieri, E., Lagoumintzis, G., Lazaridis, K., Sideri, A., Zouridakis, M., Tzartos, S.J., 2007. Muscle and neuronal nicotinic acetylcholine receptors. FEBS J. 274, 3799–3845. 10.1111/j.1742-4658.2007.05935.x

Kassambara, A., 2025. ggpubr: ‘ggplot2’ based publication ready plots. 10.32614/CRAN.package.ggpubr

Kassambara, A., 2023. rstatix: pipe-friendly framework for basic statistical tests. 10.32614/CRAN.package.rstatix

Köhler, A., Pirk, C.W.W., Nicolson, S.W., 2012. Honeybees and nectar nicotine: deterrence and reduced survival versus potential health benefits. J. Insect Physiol. 58, 286–292. 10.1016/j.jinsphys.2011.12.002

Lenth, R.V., 2025. emmeans: estimated marginal means, aka least-squares means. 10.32614/CRAN.package.emmeans

Lerch-Henning, S., Nicolson, S.W., 2013. Bird pollinators differ in their tolerance of a nectar alkaloid. J. Avian Biol. 44, 408–416. 10.1111/j.1600-048X.2013.00079.x

Lillywhite, H.B., Shine, R., Jacobson, E., Denardo, D.F., Gordon, M.S., Navas, C.A., Wang, T., Seymour, R.S., Storey, K.B., Heatwole, H., Heard, D., Brattstrom, B., Burghardt, G.M., 2017. Anesthesia and euthanasia of amphibians and reptiles used in scientific research: Should hypothermia and freezing be prohibited? BioScience 67, 53–61. 10.1093/biosci/biw143

López-Hervas, K., Santos, J.C., Ron, S.R., Betancourth-Cundar, M., Cannatella, D.C., Tarvin, R.D., 2024. Deep divergences among inconspicuously colored clades of *Epipedobates* poison frogs. Mol. Phylogenet. Evol. 195, 108065. 10.1016/j.ympev.2024.108065

Manson, J.S., Thomson, J.D., 2009. Post-ingestive effects of nectar alkaloids depend on dominance status of bumblebees. Ecol. Entomol. 34, 421–426. 10.1111/j.1365-2311.2009.01100.x

Minder, B., Brunetti, A.E., Mendonça, J.N., da Silva, R.M., Saporito, R.A., Lopes, N.P., Grant, T., 2026. The role of different organs in the hydroxylation of pumiliotoxin (+)-251D to allopumiliotoxin (+)-267A in the poison frog *Adelphobates galactonotus*. Toxicon 275, 109039. 10.1016/j.toxicon.2026.109039

Naumann, C., Hartmann, T., Ober, D., 2002. Evolutionary recruitment of a flavin-dependent monooxygenase for the detoxification of host plant-acquired pyrrolizidine alkaloids in the alkaloid-defended arctiid moth *Tyria jacobaeae*. Proc. Natl. Acad. Sci. U. S. A. 99, 6085–90. 10.1073/pnas.082674499

Neuwirth, E., 2022. RColorBrewer: ColorBrewer palettes. 10.32614/CRAN.package.RColorBrewer

O’Connell, L.A., O’Connell, J.D., Paulo, J.A., Trauger, S.A., Gygi, S.P., Murray, A.W., 2021. Rapid toxin sequestration modifies poison frog physiology. J. Exp. Biol. 224, 1–8. 10.1242/jeb.230342

R Core Team, 2025. R: a language and environment for statistical computing. R Foundation for Statistical Computing, Vienna, Austria.

Ramírez-Castañeda, V., Nixon, S., Alarcón-Naforo, D., Abderemane-Ali, F., Fitch, R., Salazar-Valenzuela, D., Minor, D., Tarvin, R., 2025. Toxin resistance mechanisms span biological scales in the Royal Ground Snake (Colubridae: Erythrolamprus reginae).

Ricciardella, L.F., Bliley, J.M., Feth, C.C., Woodley, S.K., 2010. Acute stressors increase plasma corticosterone and decrease locomotor activity in a terrestrial salamander (*Desmognathus ochrophaeus*). Physiol. Behav. 101, 81–86. 10.1016/j.physbeh.2010.04.022

Ridenhour, B.J., Brodie III, E.D., Brodie Jr, E.D., 2004. Resistance of neonates and field-collected garter snakes (*Thamnophis* spp.) to tetrodotoxin. J. Chem. Ecol. 30, 143–54.

Robinson, K.E., Moniz, H.A., Stokes, A.N., Feldman, C.R., 2024. Where does all the poison go? Investigating toxicokinetics of newt (*Taricha*) tetrodotoxin (TTX) in garter snakes (*Thamnophis*). J. Chem. Ecol. 50, 489–502. 10.1007/s10886-024-01517-7

Salehi, B., Sestito, S., Rapposelli, S., Peron, G., Calina, D., Sharifi-Rad, M., Sharopov, F., Martins, N., Sharifi-Rad, J., 2018. Epibatidine: a promising natural alkaloid in health. Biomolecules 9, 6. 10.3390/biom9010006

Santos, J.C., Tarvin, R.D., O’Connell, L.A., 2016. A review of chemical defense in poison frogs (Dendrobatidae): ecology, pharmacokinetics, and autoresistance, in: Chemical Signals in Vertebrates. pp. 305–337. 10.1007/978-0-387-73945-8

Saporito, R.A., Donnelly, M.A., Spande, T.F., Garraffo, H.M., 2012. A review of chemical ecology in poison frogs. Chemoecology 22, 159–168. 10.1007/s00049-011-0088-0

Schindelin, J., Arganda-Carreras, I., Frise, E., Kaynig, V., Longair, M., Pietzsch, T., Preibisch, S., Rueden, C., Saalfeld, S., Schmid, B., Tinevez, J.-Y., White, D.J., Hartenstein, V., Eliceiri, K., Tomancak, P., Cardona, A., 2012. Fiji: an open-source platform for biological-image analysis. Nat. Methods 9, 676–682. 10.1038/nmeth.2019

Shine, R., Amiel, J., Munn, A.J., Stewart, M., Vyssotski, A.L., Lesku, J.A., 2015. Is “cooling then freezing” a humane way to kill amphibians and reptiles. Biol. Open 4, 760–763. 10.1242/bio.012179

Soto, K.M., Hardin, F.O., Alleyne, H.P., Fischer, E.K., 2024. Individual behavioral variability across time and contexts in *Dendrobates tinctorius* poison frogs. Behav. Ecol. Sociobiol. 78, 62. 10.1007/s00265-024-03474-3

Spande, T.F., Garraffo, H.M., Edwards, M.W., Yeh, H.J.C., Pannell, L., Daly, J.W., 1992. Epibatidine: a novel (chloropyridy1)azabicycloheptane with potent analgesic activity from an Ecuadoran poison frog. J. Am. Chem. Soc. 114, 3475–3478. 10.1021/ja00035a048

Speed, M.P., Ruxton, G.D., 2014. Ecological pharmacodynamics: prey toxin evolution depends on the physiological characteristics of predators. Anim. Behav. 98, 53–67. 10.1016/j.anbehav.2014.09.011

Tadmor-Melamed, H., Markman, S., Arieli, A., Distl, M., Wink, M., Izhaki, I., 2004. Limited ability of Palestine Sunbirds *Nectarinia osea* to cope with pyridine alkaloids in nectar of Tree Tobacco *Nicotiana glauca*. Funct. Ecol. 18, 844–850. 10.1111/j.0269-8463.2004.00929.x

Tarvin, R.D., Borghese, C.M., Sachs, W., Santos, J.C., Lu, Y., O’Connell, L.A., Cannatella, D.C., Harris, R.A., Zakon, H.H., 2017a. Interacting amino acid replacements allow poison frogs to evolve epibatidine resistance. Science 357, 1261–1266. 10.1126/science.aan5061

Tarvin, R.D., Coleman, J.L., Donoso, D.A., Betancourth-Cundar, M., López-Hervas, K., Gleason, K.S., Sanders, J.R., Smith, J.M., Ron, S.R., Santos, J.C., Sedio, B.E., Cannatella, D.C., Fitch, R.W., 2024. Passive accumulation of alkaloids in inconspicuously colored frogs refines the evolutionary paradigm of acquired chemical defenses. eLife 13, RP100011. 10.7554/eLife.100011

Tarvin, R.D., Pearson, K.C., Douglas, T.E., Ramírez-Castañeda, V., Navarrete, M.J., 2023. The diverse mechanisms that animals use to resist toxins. Annu. Rev. Ecol. Evol. Syst. 54, 283–306. 10.1146/annurev-ecolsys-102320-102117

Tarvin, R.D., Powell, E.A., Santos, J.C., Ron, S.R., Cannatella, D.C., 2017b. The birth of aposematism: High phenotypic divergence and low genetic diversity in a young clade of poison frogs. Mol. Phylogenet. Evol. 109, 283–295. 10.1016/j.ympev.2016.12.035

Tarvin, R.D., Santos, J.C., O’Connell, L.A., Zakon, H.H., Cannatella, D.C., 2016. Convergent substitutions in a sodium channel suggest multiple origins of toxin resistance in poison frogs. Mol. Biol. Evol. 1–35.

Taylor, E.N., Diele-Viegas, L.M., Gangloff, E.J., Hall, J.M., Halpern, B., Massey, M.D., Rödder, D., Rollinson, N., Spears, S., Sun, B., Telemeco, R.S., 2021. The thermal ecology and physiology of reptiles and amphibians: A user’s guide. J. Exp. Zool. Part Ecol. Integr. Physiol. 335, 13–44. 10.1002/jez.2396

Taylor, E.W., Leite, C. a. C., McKenzie, D.J., Wang, T., 2010. Control of respiration in fish, amphibians and reptiles. Braz. J. Med. Biol. Res. 43, 409–424. 10.1590/S0100-879X2010007500025

Thill, V.L., Teglas, M.B., Moniz, H.A., Feldman, C.R., 2025. Ancient enemies? Patterns of Resistance to widow spider venom in lizards. Evol. Biol. 52, 120–137. 10.1007/s11692-025-09650-1

Tomar, S., 2006. Converting video formats with FFmpeg. Linux J. 2006, 10.

Wang, S.-Y., Wang, G.K., 2017. Single rat muscle Na+ channel mutation confers batrachotoxin autoresistance found in poison-dart frog *Phyllobates terribilis*. Proc. Natl. Acad. Sci. 114, 10491–10496. 10.1073/pnas.1707873114

Waters, K.R., Dugas, M.B., Grant, T., Saporito, R.A., 2023. The ability to sequester the alkaloid epibatidine is widespread among dendrobatid poison frogs. Evol. Ecol. 10.1007/s10682-023-10260-6

Wenig, P., Odermatt, J., 2010. OpenChrom: a cross-platform open source software for the mass spectrometric analysis of chromatographic data. BMC Bioinformatics 11, 405. 10.1186/1471-2105-11-405

Wickham, H., 2016. ggplot2: Elegant graphics for data analysis. Springer-Verlag New York.

Wickham, H., Averick, M., Bryan, J., Chang, W., McGowan, L.D., François, R., Grolemund, G., Hayes, A., Henry, L., Hester, J., Kuhn, M., Pedersen, T.L., Miller, E., Bache, S.M., Müller, K., Ooms, J., Robinson, D., Seidel, D.P., Spinu, V., Takahashi, K., Vaughan, D., Wilke, C., Woo, K., Yutani, H., 2019. Welcome to the tidyverse. J. Open Source Softw. 4, 1686. 10.21105/joss.01686

Williams, B.L., Brodie Jr, E.D., Brodie III, E.D., 2002. Comparisons between Toxic effects of tetrodotoxin administered orally and by intraperitoneal injection to the garter snake *Thamnophis sirtalis*. J. Herpetol. 36, 112–115. 10.2307/1565813

Yatime, L., Laursen, M., Morth, J.P., Esmann, M., Nissen, P., Fedosova, N.U., 2011. Structural insights into the high affinity binding of cardiotonic steroids to the Na+,K+-ATPase. J. Struct. Biol. 174, 296–306. 10.1016/j.jsb.2010.12.004

York, J.M., Borghese, C.M., George, A.A., Cannatella, D.C., Zakon, H.H., 2023. A potential cost of evolving epibatidine resistance in poison frogs. BMC Biol. 21, 144. 10.1186/s12915-023-01637-8

Zalucki, M.P., Brower, L.P., Alonso-M, A., 2001. Detrimental effects of latex and cardiac glycosides on survival and growth of first-instar monarch butterfly larvae *Danaus plexippus* feeding on the sandhill milkweed *Asclepias humistrata*. Ecol. Entomol. 26, 212–224. 10.1046/j.1365-2311.2001.00313.x

