## Supplemental Material 1 for "Ingestion of the potent neurotoxin epibatidine does not compromise locomotion or behavior in the poison frog *Epipedobates tricolor*"

Adriana M. Jeckel[1], Sophie J. Draper[1], Benjamin P. Deters[1], Rachel B. Weinberg[2], Neil D. Tsutsui[2], and Rebecca D. Tarvin[1]
[1] Museum of Vertebrate Zoology and Department of Integrative Biology, University of California, Berkeley, Berkeley CA 94720
[2] Department of Environmental Science, Policy, and Management, University of California, Berkeley, Berkeley CA 94720

Table SM1. Results from the generalized linear mixed models including all samples (water control, ethanol control, and epibatidine treatment samples).

| Parameter | Coefficient | X2 | d.f. | p value |
|---|---|---|---|---|
| Latency to first reaction | | | | |
| | Treatment | 7.941 | 3 | **0.047** |
| | Time | 1.758 | 2 | 0.415 |
| | Body mass | 0.000 | 1 | 0.995 |
| | Stimulation | 3.570 | 1 | 0.059 |
| | Treatment:Time | 9.637 | 6 | 0.141 |
| Average distance per jump (cm) | | | | |
| | Treatment | 11.621 | 3 | **0.009** |
| | Time | 0.119 | 2 | 0.942 |
| | Body mass | 0.158 | 1 | 0.691 |
| | Stimulation | 0.630 | 1 | 0.427 |
| | Treatment:Time | 8.107 | 6 | 0.230 |
| Jumps per second | | | | |
| | Treatment | 1.671 | 3 | 0.643 |
| | Time | 0.111 | 2 | 0.946 |
| | Body mass | 0.016 | 1 | 0.898 |
| | Stimulation | Stimulated frogs removed from analysis | | |
| | Treatment:Time | 6.827 | 6 | 0.337 |
| Average latency between jumps | | | | |
| | Treatment | 1.864 | 3 | 0.601 |
| | Time | 0.557 | 2 | 0.757 |
| | Body mass | 1.748 | 1 | 0.186 |
| | Stimulation | Stimulated frogs removed from analysis | | |
| | Treatment:Time | 7.021 | 6 | 0.319 |

Table SM2. Tukey HSD Post-Hoc analysis of latency to first reaction parameter from the generalized linear mixed models including all samples (water control, ethanol control, and epibatidine treatment samples).

| **Time & Treatment Group** | **EMM** | **95% Lower CL** | **95% Upper CL** | **SE** | **Within-Time Contrast (p-value)** | **Within-Treatment Contrast (p-value)** |
|---|---|---|---|---|---|---|
| **Time = 3 min** | | | | | | |
| Control | 8.33 | 2.62 | 26.46 | 4.91 | All comparisons p > 0.70 | All comparisons p > 0.70 |
| Treatment_2ug | 11.42 | 4.31 | 30.27 | 5.68 | | All comparisons p > 0.50 |
| Treatment_6ug | 6.10 | 2.30 | 16.16 | 3.03 | | All comparisons p > 0.70 |
| Water | 4.13 | 0.82 | 20.79 | 3.40 | | vs. 4h: Ratio = 10.61 (p = 0.0655) |
| **Time = 1 h** | | | | | | |
| Control | 6.03 | 1.34 | 27.16 | 4.63 | All comparisons p > 0.70 | All comparisons p > 0.70 |
| Treatment_2ug | 5.08 | 1.72 | 15.05 | 2.82 | | All comparisons p > 0.50 |
| Treatment_6ug | 10.68 | 3.74 | 30.51 | 5.72 | | All comparisons p > 0.70 |
| Water | 6.76 | 1.34 | 34.18 | 5.59 | | **vs. 4h: Ratio = 17.38 (p = 0.0186)** |
| **Time = 4 h** | | | | | | |
| Control | 12.88 | 3.52 | 47.17 | 8.53 | **vs. Water: Ratio = 33.13 (p = 0.0015)** | All comparisons p > 0.70 |
| Treatment_2ug | 5.35 | 1.73 | 16.59 | 3.09 | **vs. Water: Ratio = 13.76 (p = 0.0163)** | All comparisons p > 0.50 |
| Treatment_6ug | 6.54 | 2.11 | 20.26 | 3.77 | **vs. Water: Ratio = 16.83 (p = 0.0070)** | All comparisons p > 0.70 |
| Water | 0.389 | 0.09 | 1.69 | 0.291 | ***(Significantly lower than all others)*** | ***(Significantly lower than 1h)*** |

Table SM3. Tukey HSD Post-Hoc analysis of average distance per jump (cm) parameter from the generalized linear mixed models including all samples (water control, ethanol control, and epibatidine treatment samples).

| **Time & Treatment Group** | **EMM** | **95% Lower CL** | **95% Upper CL** | **SE** | **Within-Time Contrast (p-value)** | **Within-Trt Contrast (p-value)** |
|---|---|---|---|---|---|---|
| **Time = 3 min** | | | | | | |
| Control | 5.92 | 4.26 | 8.24 | 1.00 | All comparisons p > 0.20 | All comparisons p > 0.39 |
| Treatment_2ug | 6.92 | 5.37 | 8.92 | 0.90 | | All comparisons p > 0.94 |
| Treatment_6ug | 6.84 | 5.30 | 8.83 | 0.89 | | All comparisons p > 0.26 |
| Water | 9.88 | 6.56 | 14.88 | 2.06 | | All comparisons p > 0.23 |
| **Time = 1 h** | | | | | | |
| Control | 7.01 | 4.78 | 10.27 | 1.37 | All comparisons p > 0.28 | All comparisons p > 0.39 |
| Treatment_2ug | 6.88 | 5.32 | 8.89 | 0.90 | | All comparisons p > 0.94 |
| Treatment_6ug | 6.25 | 4.82 | 8.11 | 0.83 | | All comparisons p > 0.26 |
| Water | 9.63 | 6.35 | 14.59 | 2.04 | | All comparisons p > 0.23 |
| **Time = 4 h** | | | | | | |
| Control | 5.33 | 3.76 | 7.56 | 0.95 | **vs. Water: Ratio = 0.391 (p = 0.0015)**** | All comparisons p > 0.39 |
| Treatment_2ug | 7.16 | 5.37 | 9.54 | 1.05 | **vs. Water: Ratio = 0.525 (p = 0.0255)*** | All comparisons p > 0.94 |
| Treatment_6ug | 5.62 | 4.27 | 7.42 | 0.79 | **vs. Water: Ratio = 0.412 (p = 0.0006)***** | All comparisons p > 0.26 |
| Water | 13.64 | 9.45 | 19.69 | 2.56 | *(Significantly higher than all others)* | All comparisons p > 0.23 |

Table SM4. Results from the generalized linear mixed models including only epibatidine-treated samples (2-µg and 6µg treatment samples, N = 5 each) and including nanograms of epibatidine per milligram of skin mass (ng/mg) as a covariate. Due to the small sample size and lack of control samples in this model, the only covariate of interest is the effect of alkaloid (ng/mg) in the parameters.

| Parameter | Coefficient | X2 | d.f. | p value |
|---|---|---|---|---|
| Latency to first reaction | | | | |
| | Treatment | 1.304 | 1 | 0.254 |
| | Time | 2.936 | 3 | 0.402 |
| | **Alkaloid_ng_mg** | **0.807** | **1** | **0.369** |
| | Body mass | 1.777 | 1 | 0.183 |
| | Stimulation | 1.291 | 1 | 0.256 |
| | Treatment:Time | 4.577 | 3 | 0.206 |
| Average distance per jump (cm) | | | | |
| | Treatment | 1.033 | 1 | 0.309 |
| | Time | 3.205 | 3 | 0.361 |
| | **Alkaloid_ng_mg** | **0.812** | **1** | **0.368** |
| | Body mass | 0.000 | 1 | 0.989 |
| | Stimulation | 1.602 | 1 | 0.206 |
| | Treatment:Time | 20.971 | 3 | 0.000 |
| Jumps per second | | | | |
| | Treatment | 0.000 | 1 | 1.000 |
| | Time | 7.042 | 3 | 0.071 |
| | **Alkaloid_ng_mg** | **0.042** | **1** | **0.837** |
| | Body mass | 0.963 | 1 | 0.326 |
| | Stimulation | Stimulated frogs removed from analysis | | |
| | Treatment:Time | 8.372 | 3 | 0.039 |
| Average latency between jumps | | | | |
| | Treatment | 0.100 | 1 | 0.751 |
| | Time | 20.387 | 3 | 0.000 |
| | **Alkaloid_ng_mg** | **1.066** | **1** | **0.302** |
| | Body mass | 1.442 | 1 | 0.230 |
| | Stimulation | Stimulated frogs removed from analysis | | |
| | Treatment:Time | 8.713 | 3 | 0.033 |
